# Attention Modulates Stimulus Representations in Neural Feature Dimension Maps

**DOI:** 10.1101/2025.01.10.632497

**Authors:** Daniel D. Thayer, Thomas C. Sprague

**Affiliations:** Department of Psychological and Brain Sciences, University of California, Santa Barbara

**Keywords:** feature-selective attention, visual system, fMRI, priority map

## Abstract

Computational theories posit that attention is guided by a combination of spatial maps for individual features that can be dynamically weighted according to task goals. Consistent with this framework, when a stimulus contains several features, attending to one or another feature results in stronger fMRI responses in regions preferring the attended feature. We hypothesized that multivariate activation patterns across feature-responsive cortical regions form spatial ‘feature dimension maps’, which combine to guide attentional priority. We tested this prediction by reconstructing spatial maps from fMRI activation patterns across retinotopic regions of visual cortex while participants performed a feature-selective attention task. Participants viewed a peripheral visual stimulus at a random location which always contained moving colored dots. On each trial, participants were precued to report the predominant direction of motion or color of the stimulus, or to attend fixation. Stimulus representations in reconstructed maps based on a spatial inverted encoding model were selectively enhanced in color-selective regions when color was attended, and in motion-selective regions when motion was attended. While enhancement was localized to the stimulus position in color-selective regions, modulations in motion-selective regions was consistent with a more global enhancement when motion was task relevant. These results suggest feature-selective cortical regions support ‘neural feature dimension maps’: spatial maps of different visual features that are dynamically reweighted based on task demands to guide visual behavior to the most relevant locations based on important features

## Introduction

The environment typically contains many objects competing for our attention. While gardening, your goal may be to find green weeds in a strawberry bush, so attention will be directed to green objects. However, task-irrelevant objects that are salient may capture your attention, such as a hummingbird darting in the sky or a red strawberry. These two factors contributing to how attention is allocated in the visual field—task-relevant goals and task-irrelevant salience—are core components of priority map theory (Awh et al., 2012; Itti & Koch, 2001; Luck et al., 2020; Serences & Yantis, 2006; Treisman & Gelade, 1980; Wolfe, 1994). Per this theory, these priority signals are integrated to index the most important locations in a scene, and locations with the highest level of priority are then selected for further processing (Carrasco, 2011; Eckstein, 2011).

In this framework, overall priority is computed based on the combination of priority indexed independently for individual feature dimensions, like color, motion, orientation, and shape. Priority within these ‘feature dimension maps’ is a combination of salience (defined based on image-computable properties, like local feature contrast) and relevance (defined based on task goals, like which location(s) or feature(s) are important; Wolfe & Horowitz, 2004, 2017). For example, a color dimension map would index locations made salient based on their unique color relative to other stimuli, while a motion dimension map would do the same for stimuli moving in a unique direction. Additionally, it would be expected that making one feature task-relevant would sculpt stimulus representations in the corresponding map, leading to a greater impact on downstream priority maps (Bahle et al., 2019; Corbetta & Shulman, 2002; Folk et al., 1992; Leber & Egeth, 2006; Moore & Armstrong, 2003; Poltoratski et al., 2017; Witkowski & Geng, 2023).

Recently, we demonstrated that color- (hV4/VO1/VO2; Bartels & Zeki, 2000; Brewer et al., 2005) and motion- (TO1/TO2; Amano et al., 2009; Huk et al., 2002) selective visual cortical regions index salient locations on the screen based on their preferred feature values, even when those locations are task-irrelevant (Thayer & Sprague, 2023). Importantly, studies investigating the impact of task relevance on feature-selective regions cannot establish whether feature-selective regions satisfy predictions of neural feature dimension maps because they all either measured from a single feature-selective visual region (Bichot et al., 2005; Martínez-Trujillo & Treue, 2002; Mazer & Gallant, 2003; Ogawa & Komatsu, 2004, 2006; Reynolds & Desimone, 2003), include only a single goal-relevant feature dimension (Beck & Kastner, 2005; Cook & Maunsell, 2002; Moran & Desimone, 1985), or investigate goal-driven prioritization of a stimulus location without selective attention to specific features (Bisley & Goldberg, 2003, 2006; Ikkai & Curtis, 2008; Sprague, Itthipuripat, et al., 2018; Sprague & Serences, 2013). If the same regions selectively encode representations of salient locations based on their preferred feature value and reweight their representations based on an observer’s task demands, this would provide further support for a role of compartmentalized ‘neural feature dimension maps’ in computing attentional priority.

Task demands can sculpt neural responses in visual cortex through spatial attention (Desimone & Duncan, 1995; Kastner et al., 1999; Sprague, Itthipuripat, et al., 2018) and feature-selective attention (Chawla et al., 1999; Saenz et al., 2002; Serences & Boynton, 2007; Treue & Trujillo, 1999). Spatial attention improves neural representation of an attended location relative to other irrelevant locations in the visual field as shown by changes in fMRI BOLD responses (Ikkai & Curtis, 2008; Kastner et al., 1999, 2001; Sprague, Itthipuripat, et al., 2018; Sprague & Serences, 2013; Vo et al., 2017). Attention to specific feature values has instead been shown to have a spatially global effect, where all instances of a goal-relevant feature value, such as the color green, are prioritized irrespective of stimulus location or relevance. Previous work has demonstrated that feature-selective attention modulates fMRI BOLD activity in color- and motion-selective regions (Beauchamp et al., 1997; Chawla et al., 1999; McMains et al., 2007; O’Craven et al., 1999; Runeson et al., 2013; Saenz et al., 2002), by enhancing responses corresponding to stimuli matching the attended feature value across the entire screen, even at locations containing no stimulus (Serences & Boynton, 2007). Since both relevance signals can together sculpt neural response patterns within feature-selective regions of cortex, it is important to characterize their joint influence on activation profiles supporting neural feature dimension maps.

While both relevance of a stimulus location (spatial attention) and relevance of a specific feature dimension (feature-selective attention) have each been shown to impact activation levels at stimulus locations in feature-selective regions, it is unknown whether feature-selective modulations occur equivalently across the entire visual field, reflecting a *global enhancement* of processing of the attended feature dimension across the entire map, or instead, are primarily restricted to the location of the presented stimuli containing the attended feature dimension, reflecting a *local enhancement* of relevant locations within the attended map (Fig. 1; McMains et al., 2007).

**Figure 1:**
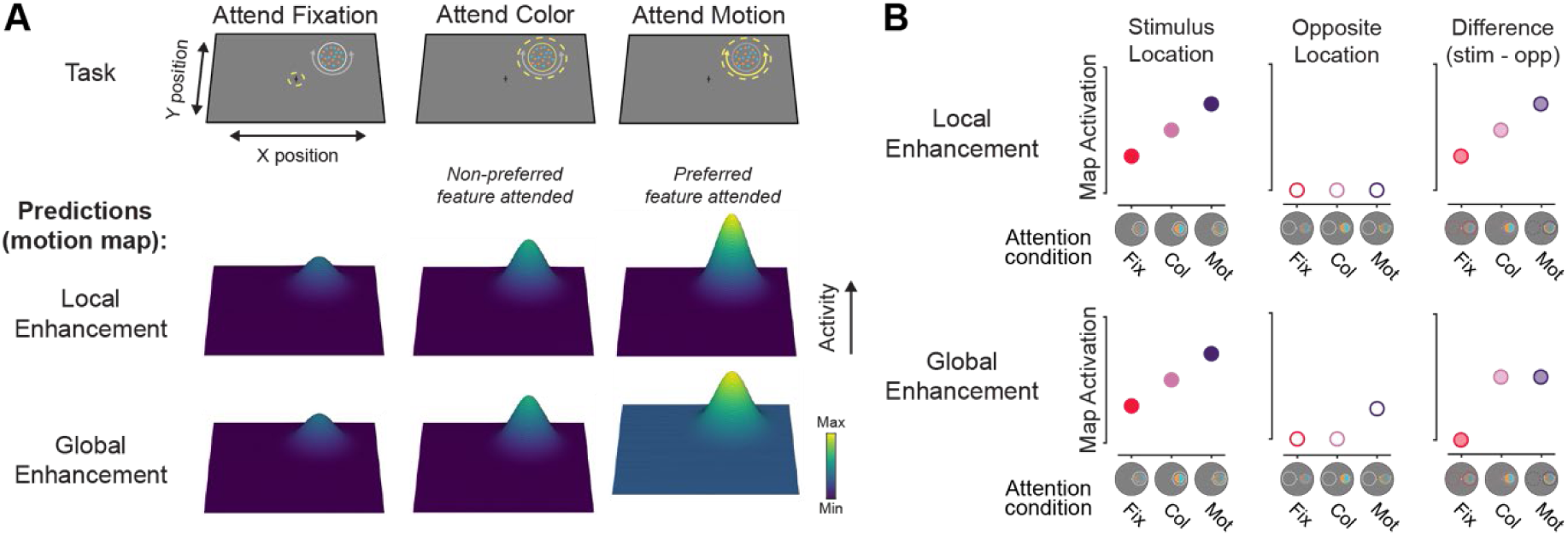
How does attention to a specific feature dimension sculpt neural feature dimension maps? **A.** Priority map theory posits that various “feature dimension maps” are used to index the most important locations in the visual field based on computations within their preferred feature dimension, and that activation in these maps should scale with an observer’s goals. If detecting or discriminating motion is needed for an ongoing task (e.g., identifying the motion direction of a darting hummingbird), then activation within the corresponding “motion map” would increase the importance associated with the hummingbird’s location. There are two ways in which the motion map could prioritize information beyond the local effects expected of spatial attention (e.g., Sprague et al, 2018). Local enhancement could occur, such that only the location of a stimulus with the attended feature is prioritized. Alternatively, global enhancement could occur, such that activation across the entire map is additively scaled, leading to increased sensitivity to the attended feature dimension at any location. This type of modulation would still drive a stronger target representation but would additionally lead to stronger responses in locations that contain no stimulus when motion is the goal-relevant feature dimension. A motion dimension map is depicted here, but modulations would similarly apply for other feature dimensions, such as color. **B.** Evaluating the activation at the stimulus location and the opposite location within the feature (motion) map can discriminate between the local and global enhancement accounts. Both models predict that the stimulus will have the greatest activation at the stimulus location when the preferred feature dimension is relevant (e.g., motion; left). If enhancement is localized, then activation at the opposite location should not change across conditions (center). However, if there is global enhancement, then the opposite location should have increased activation when motion is task relevant. By computing the difference in activation between the stimulus and opposite locations, the spatial specificity of feature-selective modulations can be assessed (right). If this activation difference (stimulus-opposite) is greater for the attend motion condition in the motion map, enhancement is localized. However, if the activation difference is similar between the attend color and motion conditions, then enhancement is global across the feature dimension map.

Here, we used a multivariate inverted encoding model (IEM; Thayer & Sprague, 2023) approach to determine how attention to a cued feature dimension modulated activity across neural feature dimension maps encoding an identical visual stimulus. We found that map locations corresponding to the stimulus location in motion- and color-selective regions had the greatest activation, and that these stimulus representations were strongest when attending to each region’s preferred feature. Interestingly, feature-specific modulations in color regions showed evidence for local enhancement while modulations in motion regions more closely resembled global enhancement, suggesting that each feature dimension map was uniquely modulated by feature-selective attention. These findings were only observable after implementing IEM and leveraging region-level spatial activation patterns, demonstrating the necessity to consider the information jointly encoded across entire neural populations when characterizing the computational properties of neural feature dimension maps.

## Materials & Methods

### Participants

We recruited 10 subjects from the University of California, Santa Barbara (UCSB) community to participate in this study (9 female, 18-29 years old). We identified an appropriate sample size for our main effect of interest by conducting a power analysis using pilot data (n = 3). In the pilot study, we had a strong effect (d_z_ = 1.24) which required a sample of 6 subjects to obtain 80% power with an alpha criterion of α = .05 as indicated by a power analysis conducted in G*Power (Faul et al., 2007). We collected a large number of measurements from each subject to minimize within-subject variance, which can benefit statistical power more than increased sample sizes (Baker et al., 2021). Subjects reported normal or corrected-to-normal vision and did not report neurological conditions. All procedures were approved by the UCSB Institutional Review Board (IRB# 5-24-0030), and the study was registered on ClinicalTrials.gov (NCT06281457). Subjects gave written informed consent before participating and were compensated for their time ($20/h for scanning sessions, $10-20/h for behavioral familiarization/training).

### Stimuli and Procedure

Participants performed a 30-minute training session before scanning so that they were familiarized with the instructions. We used this session to establish the initial behavioral performance thresholds used in the first run of the scanning session. In the main task session, participants were scanned for a single two-hour session consisting of at least 4 runs of a spatial mapping task, which were used to independently estimate encoding models for each voxel, and 8 runs of the experimental feature-selective attention task. All participants also underwent additional anatomical and retinotopic mapping scanning sessions (1-2x 1.5-2 hr sessions) to identify regions of interest (ROI; see Region of interest definition).

Stimuli were presented using the Psychophysics toolbox (Brainard, 1997; Pelli, 1997) for MATLAB (The MathWorks, Natick, MA). Visual stimuli were rear-projected onto a screen placed ∼110 cm from the participant’s eyes at the head of the scanner bore using a contrast-linearized LCD projector (1,920×1,080, 60 Hz) during the scanning session. For two of the participants, a CRS BOLDScreen LCD monitor was used to present stimuli (1,920×1,080, 60 Hz) that was placed ∼149 cm from the participant’s eyes. Participants who performed the task on the BOLDScreen had slightly different stimulus dimensions (scaled by a factor of .9 compared to the projector), but all results are qualitatively identical for the subjects who used either the BOLDScreen monitor or the projector, and all findings hold considering only the n = 8 participants tested with the projector. Throughout, we describe stimulus dimensions/units based on the projector stimulus size. In the behavioral familiarization session, we presented stimuli on a contrast-linearized LCD monitor (2,560×1,440, 60 Hz) 62 cm from participants, who were seated in a dimmed room and positioned using a chin rest. For all sessions and tasks (main tasks, localizers, and mapping task), we presented stimuli on a neutral gray circular aperture (9.15° radius), surrounded by black (only aperture shown in Fig. 2).

**Figure 2:**
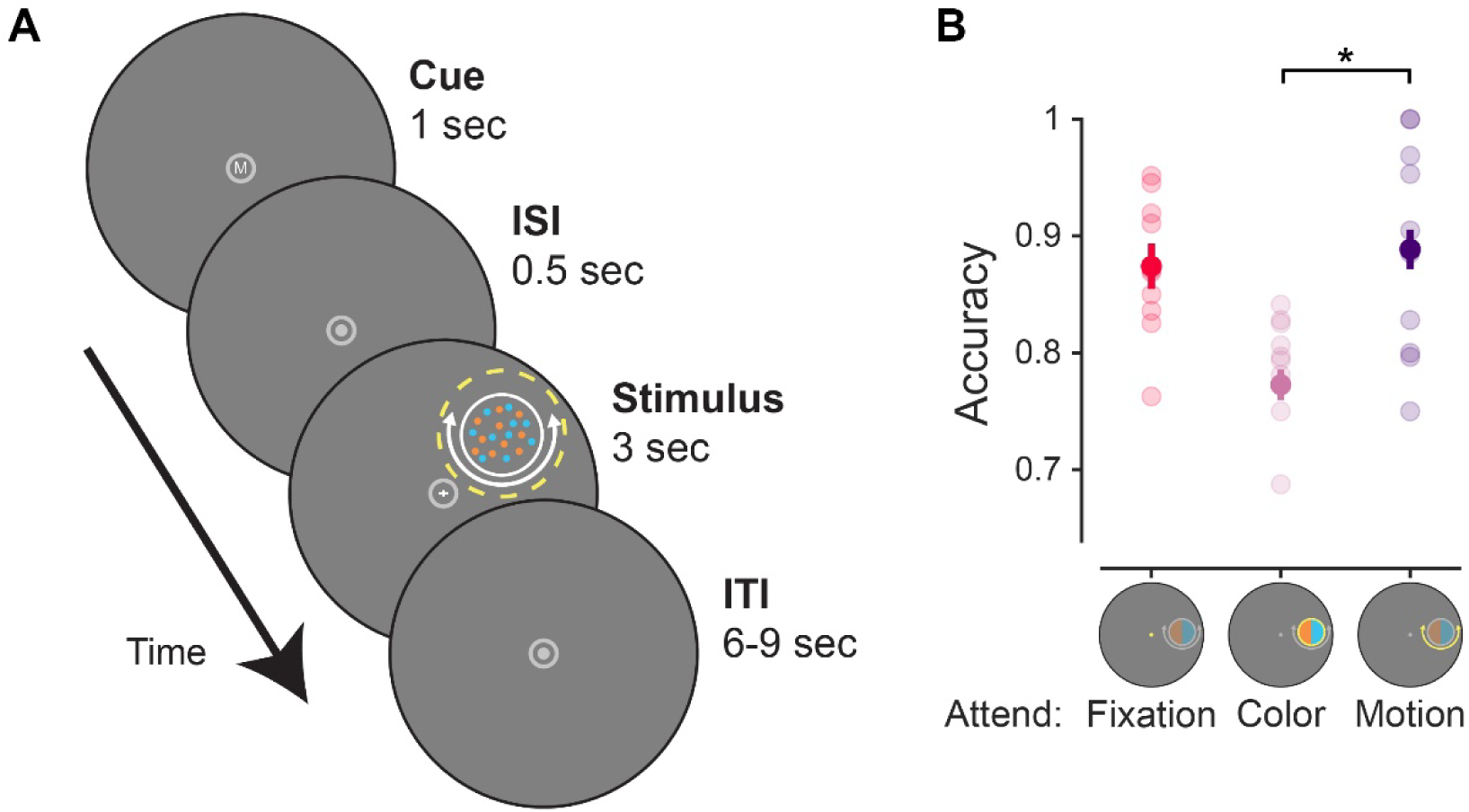
Feature-selective attention task. **A.** On each trial, participants viewed a dot array comprised of colorful moving dots. All dots were randomly either cyan or orange, and all dots were always moving, but only some would move clockwise or counterclockwise, while the rest moved in random direction. A flickering fixation cross (3 Hz) was also always present, and the vertical or horizontal arm of the cross would increase in size once during the stimulus presentation period. A letter at fixation cued participants to perform one of three attention tasks related to the upcoming stimulus display (F: attend fixation, C/M attend peripheral stimulus color/motion, respectively). On attend fixation trials, participants reported whether the horizontal or vertical arm of the fixation cross lengthened; on attend color/motion trials, they reported the prominent feature value of the cued feature dimension (more cyan/orange dots, or more dots moving clockwise/counterclockwise). **B.** Accuracy for each attention condition. Performance was slightly worse for the attend color than the attend motion or fixation conditions. Error bars indicate within-subject SEM (Franz & Loftus, 2012). Faint dots indicate individual subject means. * Significant difference based on permuted paired-samples *t* test (*p* < 0.05).

**Figure 3:**
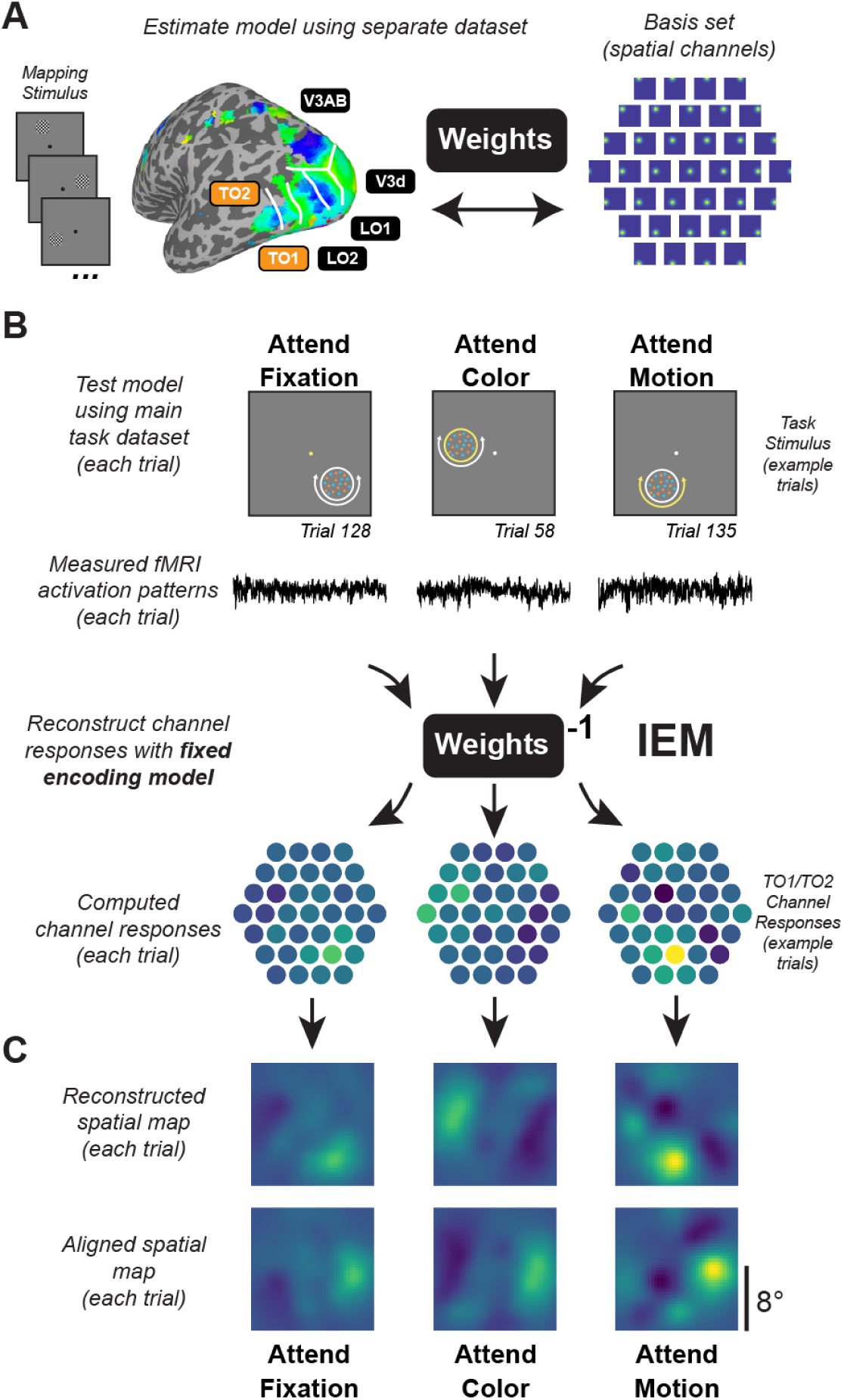
Inverted encoding model used to reconstruct maps of the visual field. **A.** Data from an independent spatial mapping task was used to estimate a spatial IEM for each ROI (for details, see Materials and Methods). **B.** Using this approach, activation patterns across all voxels in a region for each trial can be mapped onto the activation of spatial channels that are then summed to produce reconstructed spatial maps of the visual field. Identical encoding models, estimated with the task in (A), are used to compute reconstructions for each condition, ensuring results can be compared on equal footing. We reconstructed maps for each trial using activation patterns averaged 4.5-7.5 s after cue onset and directly compared the map activation across conditions. **C.** Since maps are in visual field coordinates, we were able to rotate and align reconstructed spatial maps to the known position of the moving dot array, then average across trials within each condition. Computed channel responses are example data from a representative subject’s (sub015) aggregate TO1/TO2 ROI.

### Feature-selective attention task

For each trial of the main task (Fig. 2), participants viewed a colorful moving dot stimulus in the periphery (similar to that used in Mante et al., 2013) along with a flickering central fixation cross (0.2° size). The colorful moving dot stimulus was a 1.5°-radius circular disc comprised of several individual dots centered 5° from fixation at a random location along an invisible ring. Individual dots were physically isoluminant, colored either cyan (RGB: [0; 200; 200]) or orange (RGB: [255; 165; 0]), and moved clockwise or counterclockwise around the stimulus center, or in a random direction. Individual dots occupied 0.04° of visual angle, with a dot density was 40 dots/deg^2^, and moved at a speed of 4.5°/s. Dots were randomly replotted every 50 ms or when they exceeded the stimulus bounds. Both cyan and orange dots were always present. Dots were always moving, but only some dots on each trial would move clockwise or counterclockwise, while the rest of the dots moved in a random direction (uniform distribution of planar motion directions).

At the start of each trial, participants were shown a letter cue at fixation (0.4° height) that would indicate what aspect of the display to attend. The letter cue was presented for 1000 ms in white (RGB: [180; 180; 180]) Arial font. The cue could be an ‘F’ (attend fixation), ‘C’ (attend color), or ‘M’ (attend motion). After a 500 ms blank period, with only the aperture and fixation dot present, the colorful moving dot stimulus and fixation cross was shown for 3000 ms. When cued to attend the color of the stimulus (‘C’), participants reported which individual feature value (cyan or orange) was most prevalent in the stimulus. When cued to attend the motion of the stimulus (‘M’), they reported the coherent motion direction of the stimulus (clockwise or counterclockwise). Since motion and color was present on each trial, visual input was constant, and we were able to isolate changes in the neural stimulus representations across brain regions due to instructed task goals. We adjusted the difficulty of the attend color/motion tasks between runs by changing the coherence of the dots to try and equate task performance between conditions (e.g., 80% dot coherence indicates that 80% of the dots were presented in cyan or were moving clockwise). The first run of the task in the scanner used coherence values acquired from the behavioral training session.

As a neutral baseline condition, participants attended the central fixation cross on some trials. The vertical and horizontal lines of the fixation cross were 0.2° of visual angle long (0.1° from center in either direction) and flickered at 3 Hz (10 frames on, 10 frames off at 60 Hz). On each trial, one line of the fixation cross would slightly change in size for a single flicker cycle. When cued to attend fixation (‘F’), participants reported which line increased in length (horizontal or vertical) with a button press. Since the dot stimulus was ignored on these trials due to fixation task difficulty (Thayer & Sprague, 2023), the neural response associated with the stimulus on these trials was purely driven by visual input. We adjusted the difficulty of the fixation task between runs by altering the degree of size change for vertical/horizontal lines based on behavioral accuracy (range: 0.21° to 0.32°). The fixation target appeared at a random time during the stimulus presentation period other than the first or last 500 ms.

Participants completed 8 runs of the task, where each run contained 24 trials, with 8 trials of each condition (attend fixation, attend stimulus color, attend stimulus motion). All trials were separated by a randomly selected ITI ranging from 6-9 s with an average ITI of 7.5 s. Trial order was shuffled within run. Each run started with a 3 s blank period and ended with a 10.5 s blank period, for a total run duration of 301.5 s. Eye position (right eye) was monitored throughout the experiment using an Eyelink 1000 eyetracker (SR Research) at 500 Hz. Participants performed a 5-point calibration at the start of the experiment.

### Spatial mapping task

We also acquired several runs of a spatial mapping task used to independently estimate a spatial encoding model for each voxel, following previous studies (Sprague et al., 2016; Sprague, Itthipuripat, et al., 2018; Thayer & Sprague, 2023). On each trial of the mapping task, we presented a flickering checkerboard at different positions selected from a hexagonal grid spanning the screen. Participants viewed these stimuli and responded whenever a rare contrast change occurred (6 out of 43 trials, 13.9%), evenly split between contrast increments and decrements.

The checkerboard stimulus was the same size as the stimulus in the feature-selective attention task (1.5° radius) and was presented at 70% contrast and 6-Hz full-field flicker. All stimuli appeared within a gray circular aperture with a 9.15° radius, as in the feature-selective attention task. For each trial, the location of the stimulus was selected from a hexagonal grid of 37 possible locations with an added random uniform circular jitter (0.5° radius). The base position of the grid was rotated by 30° on every other scanner run to increase spatial sampling density. As a result, every mapping trial was unique, which enabled robust spatial encoding model estimation.

Each trial started with a 3000 ms stimulus presentation period. If a target was present, then the stimulus would be dimmed/brightened for 500 ms with the stipulation that the contrast change would not occur in either the first or last 500 ms of the stimulus presentation period. Finally, ITIs were randomly selected to be either 6 or 8.25 s. All target-present trials were discarded when estimating the spatial encoding model. Each run consisted of 43 trials (6 of which included targets). We also included a 3 s blank period at the beginning of the run and a 10.5-s blank period at the end of the run. Each run totaled 432 s.

### Retinotopic mapping task

We used a previously reported task (Mackey et al., 2017; Thayer & Sprague, 2023) to identify retinotopic regions of interest (ROIs) via the voxel receptive field (vRF) method (Dumoulin & Wandell, 2008). Each run of the retinotopy task required participants attend several random dot kinematograms (RDK) within bars that would sweep across the visual field in 2.25 s steps. Three equally sized bars were presented on each step and the participants reported in which of the two peripheral bars the motion in the central bar matched with a button press. Participants received feedback via a red (incorrect) or green (correct) color change at fixation. We used a three-down/one-up staircase to maintain ∼80% accuracy throughout each run so that participants would continue to attend the RDK bars. RDK bars swept 17.5° (15.8° on BOLDScreen) of the visual field. Bar width and sweep direction was pseudo-randomly selected from several different widths (ranging from 2.0° to 7.5°) and four directions (left-to-right, right-to-left, bottom-to-top, and top-to-bottom).

### fMRI acquisition and MRI preprocessing

All scanning was conducted at the UCSB Brain Imaging Center with a 3T Siemens Prisma scanner. fMRI data acquisition and preprocessing pipelines in the current study exactly follow our previous report (Thayer & Sprague, 2023). All fMRI scans for retinotopic mapping, model estimation, and main experimental task were acquired using the CMRR MultiBand accelerated EPI pulse sequences. We acquired all images with the Siemens 64 channel head/neck coil with all elements enabled. We collected 3 T1-weighted and 1 T2-weighted anatomical images using the Siemens product MPRAGE and Turbo Spin-Echo sequences (0.8 mm isotropic voxels, 256 X 240 mm slice FOV). The TE/TR of each anatomical image were: 2.24/2400 ms (T1w) and 564/3200 ms (T2w). These scans were used to coregister all functional images to each participant’s native anatomic space. We acquired functional images using a Multiband (MB) 2D GE-EPI scanning sequence with MB factor of 4, acquiring 44 2.5 mm interleaved slides with no gap, with isotropic voxel size being 2.5 mm. The TE/TR of these scans were: 30/750 ms (4x simultaneous-multi-slice acceleration). We used freesurfer’s recon-all script (version 6.0) to process all anatomical scans and AFNI’s afni_proc.py command to preprocess all functional scans. Preprocessing included unwarping, alignment, rigid-body motion correction (6-parameter affine transform), and smoothing along the cortical surface normal. Retinotopic mapping data was spatially smoothed along the cortical surface (5 mm FWHM) for identifying ROIs, but feature-selective attention and spatial mapping task data was not smoothed beyond projecting to surface space and returning to volume space.

### Region of interest definition

We identified 15 ROIs using independent retinotopic mapping data. We fit a vRF model for each voxel in the cortical surface (in volume space) using averaged and spatially smoothed (on the cortical surface; 5 mm FWHM) time series data across all retinotopy runs (8-12 per participant). We used a compressive spatial summation isotropic Gaussian model (Kay et al., 2013; Mackey et al., 2017) as implemented in a customized, GPU-optimized version of mrVista (see Mackey et al., 2017) for detailed description of the model). High-resolution stimulus masks were created (270 x 270 pixels) to ensure similar predicted responses within each bar size across all visual field positions. Model fitting began with an initial high-density grid search, followed by subsequent nonlinear optimization. Retinotopic ROIs (V1, V2, V3, V3AB, hV4, LO1, LO2, VO1, VO2, TO1, TO2, IPS0-3) were then delineated by projecting vRF best-fit polar angle and eccentricity parameters with variance explained ≥5% onto each participant’s inflated cortical surfaces via AFNI and SUMA. We drew ROIs on each hemisphere’s cortical surface based on previously-established polar angle reversal and foveal representation criteria (Amano et al., 2009; Mackey et al., 2017; Swisher et al., 2007; Wandell et al., 2007; Winawer & Witthoft, 2015). Finally, ROIs were projected back into volume space to select voxels for analysis. For primary analyses (Figs. 4, 6), we focused on previously identified ‘feature dimension maps’ (Thayer & Sprague, 2023) which were aggregate ROIs made up of a combination of all voxels across retinotopic maps with similar feature selectivity (color: hV4, VO1, and VO2; motion: TO1 and TO2). For completeness, we additionally report results from each individual ROI (Figs. 5, 7).

**Figure 4:**
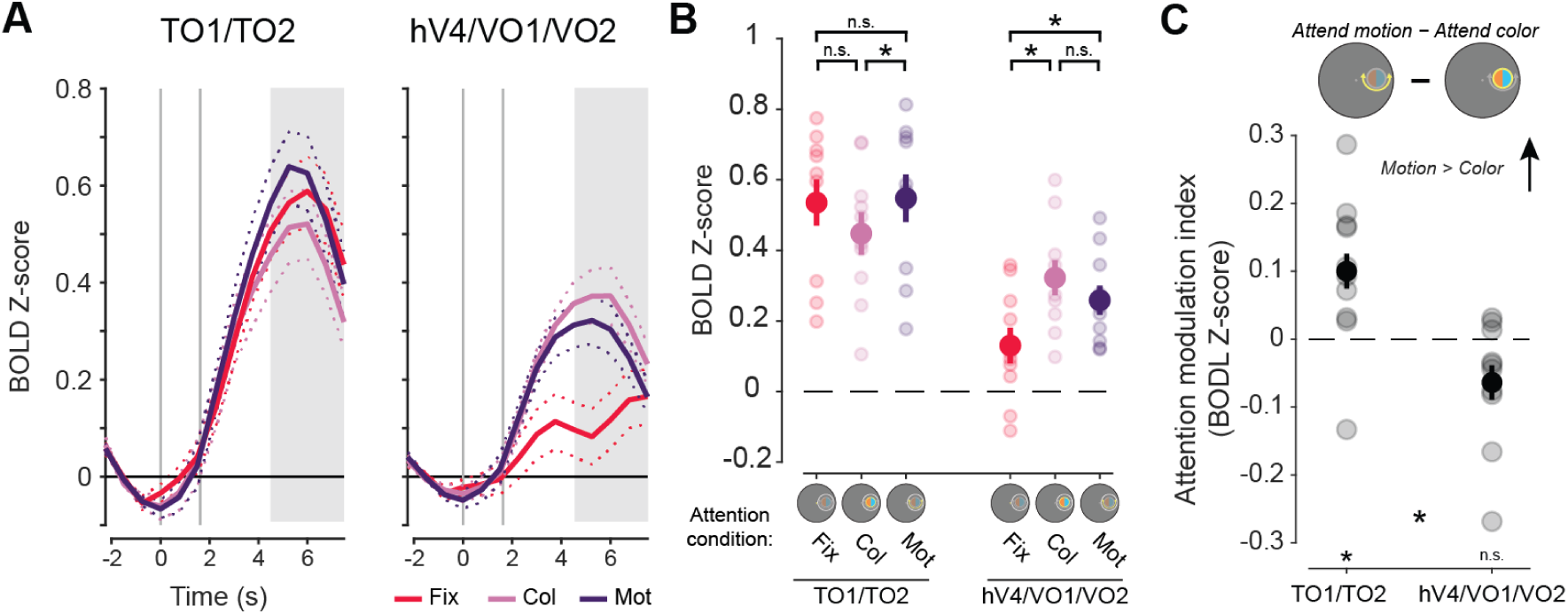
BOLD response across all voxels in aggregate feature-selective visual cortex ROIs show motion selectivity for TO1/TO2 and task demand modulation in hV4/VO1/VO2. **A.** HRFs from aggregate feature-selective retinotopic regions TO1/TO2 and hV4/VO1/VO2. Gray lines indicate cue (first) and stimulus (second) onsets. Shaded area indicates time points used for trial averaging across participants. Dashed lines indicate SEM within participants. Dot array stimulus was presented 1.5 to 4.5 seconds (second gray line to start of shaded area). **B.** Mean BOLD response across participants for each attention condition. Motion-selective regions showed no modulation of response when attending to either feature of the stimulus (color or motion) relative to when attending to fixation (color vs fixation: *p* = 0.07; motion vs fixation: *p* = 0.59). Color-selective regions were modulated by the task-relevance of the peripheral stimulus, with a greater region-level response when attending to the color (*p* < 0.01) or the motion (*p* < 0.01) relative to attending to fixation. **C.** Attention modulation index for both aggregate feature-selective regions averaged across participants. Only motion-selective regions had a response that differed from zero (*p* = 0.02). * indicate significant differences based on permuted paired or single sample *t* tests (*p* < 0.05). Error bars indicate within-subject SEM. Faint dots indicate individual subject means. Col: color, Mot: motion, Fix: fixation.

**Figure 5:**
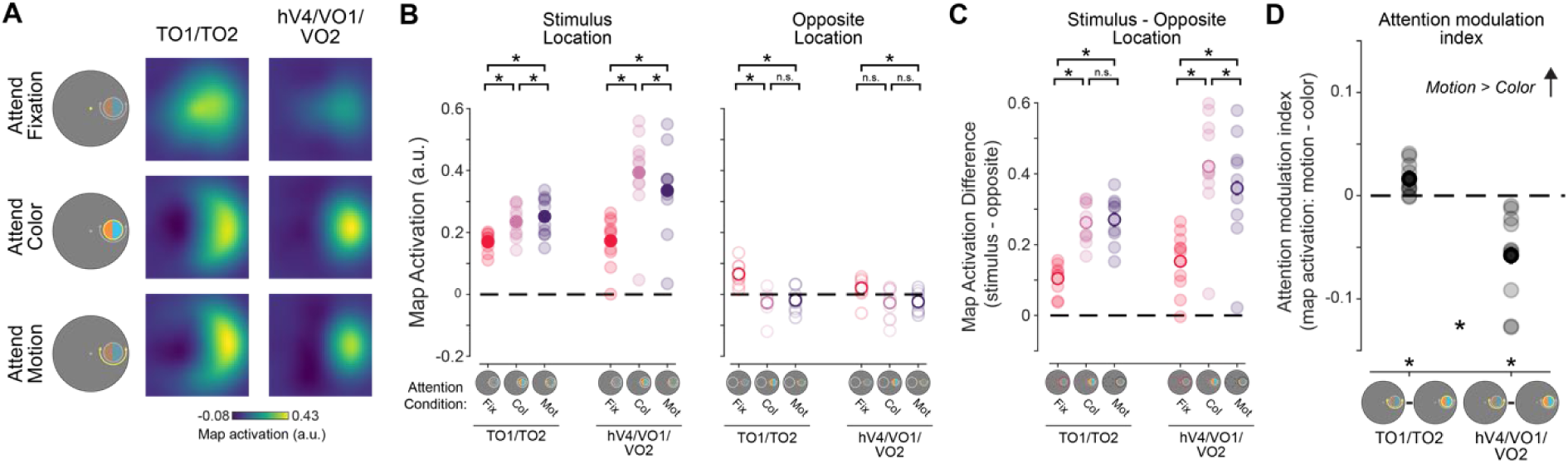
Reconstructed spatial maps from color-and motion-selective regions are differentially modulated by feature-selective attention. **A.** Reconstructions from motion- and color-selective regions from the attend fixation, color, and motion conditions. Qualitatively, there is a strong response at the aligned location, particularly when attending to either the motion or color of the stimulus. **B.** We determined whether each region was modulated by task demands by assessing the strength of the stimulus representation for each stimulus condition as well as the opposite location. For both regions, the neural response at the stimulus location was stronger when attending to either feature of the stimulus as compared to attend fixation trials, indicating that stimulus representations in these regions track task-relevance of the stimulus location. At the opposite location, responses were stronger when attending to fixation. **C.** Difference in map activation between the stimulus and opposite locations for each stimulus condition, which indexes spatial specificity of map activation profiles. Responses were always greater when attending to the stimulus as compared to fixation for both regions. While color regions hV4/VO1/VO2 show a difference between the attend motion and color conditions (*p* = 0.02), indicating local enhancement, motion regions TO1/TO2 showed no difference (*p* = 0.61), indicating global enhancement. **D.** Difference in map activation at the stimulus location between the attend motion and attend color conditions. Responses reliably differed from regions had a stronger response when attending to color. Responses differed between regions. * Significant difference based on permuted paired-samples *t* test (*p* < 0.05). Faint dots indicate individual subject means. Error bars indicate within-subject SEM.

### Inverted encoding model

We used a spatial inverted encoding model (IEM) to reconstruct images of stimulus-related activation patterns measured across entire ROIs (Sprague & Serences, 2013; Fig. 3). To do this, we first estimated an encoding model, which describes the sensitivity profile over the relevant feature dimension for each voxel in a region. This requires using data set aside for this purpose, referred to as the “training set”. Here, we used data from the spatial mapping task as the independent training set. The encoding model across all voxels within a given region is then inverted to estimate a mapping used to transform novel activation patterns from a “test set” (runs from the feature-selective attention task) and reconstruct the spatial representation of the stimulus at each timepoint. For this analysis, we used voxels for which the best-fit vRF model explained ≥5% of the variance during the retinotopic mapping task.

We built an encoding model for spatial position based on a linear combination of 37 spatial filters (Sprague, Itthipuripat, et al., 2018; Thayer & Sprague, 2023). Each voxel’s response was modeled as a weighted sum of each identically shaped spatial filter arrayed in a triangular grid (Fig. 3A). The centers of each filter were spaced by 2.83° and were Cosine functions raised to the 7^th^ power:

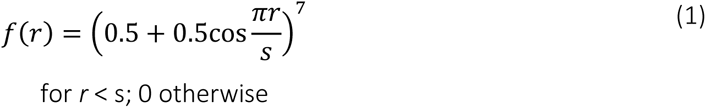

where *r* is the distance from the filter center and *s* is a size constant. The size constant reflects the distance from the center of each spatial filter at which the filter returns to 0. This triangular grid of filters forms the basis set, or information channels for our analysis. For each stimulus used in our mapping task, we converted from a contrast mask to a set of filter activation levels by taking the dot product of the vectorized stimulus mask (*n* trials × *p* pixels) and the sensitivity profile of each filter (*p* pixels × *k* channels). We then normalized the estimated filter activation levels such that the maximum activation was 1 and used this output as *C1* in the following equation, which acts as the forward model of our measured fMRI signals:

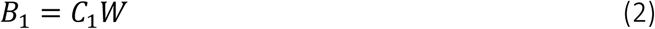

*B_1_* (*n* trials × *m* voxels) in this equation is the measured fMRI activity of each voxel during the visuospatial mapping task and *W* is a weight matrix (*k* channels × *m* voxels) which quantifies the contribution of each information channel to each voxel. *Ŵ* can be estimated using ordinary least-squares linear regression to find the weights that minimize the differences between predicted values of *B* and the observed *B_1_*:

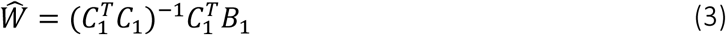

This is computed for each voxel within a region independently, making this step univariate. The resulting *Ŵ* represents all estimated voxel sensitivity profiles. We then used *Ŵ* and the measured fMRI activity of each voxel (i.e., BOLD response) during each trial (using each TR from each trial, in turn) of the feature-selective attention task using the following equation:

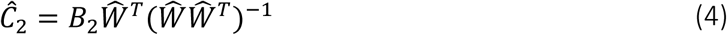

Here, *Ĉ*_2_represents the estimated activation of each information channel (*n* trials × *k* channels) which gave rise to that observed activation pattern across all voxels within a given ROI (*B_2_*; *n* trials × *m* voxels). To aid with visualization, quantification, and coregistration of trials across stimulus positions, we computed spatial reconstructions using the output of Equation 4. To do this, we weighted each filter’s spatial profile by the corresponding channel’s reconstructed activation level and then sum all weighted filters together (Fig. 3B).

Since stimuli in the feature-selective attention task were randomly positioned on every trial, we rotated the center position of spatial filters such that the resulting 2D reconstructions of the stimuli were aligned across trials and participants (Fig. 3C). We then sorted trials based on condition (attend fixation, attend color, attend motion). Finally, we averaged the 2D reconstructions across trials within the same condition for individual participants, then across all participants for our grand-average spatial reconstructions (Fig. 5; Appendix Fig. A2). Individual values within the 2D reconstructed spatial maps correspond to map activation at the corresponding relative visual field coordinates based on activation profiles across entire ROIs.

Critically, because we reconstructed all trials from all conditions of the feature-selective attention task using an identical spatial encoding model estimated with an independent spatial mapping task, we can compare reconstructions across conditions on the same footing (Sprague, Adam, et al., 2018; Sprague et al., 2019; Sprague, Itthipuripat, et al., 2018).

### Quantifying neural responses

First, to establish the impact of feature-selective attention to specific stimulus dimensions on overall activation in each ROI at the broadest spatial scale, we computed the average activation across all visually responsive voxels (defined as those with at least 5% variance explained based on vRF model fits) in each ROI (McMains et al., 2007; Runeson et al., 2013). Second, because we can measure activation across each entire region simultaneously (that is, voxels with the full spread of spatial encoding properties), we performed a multivariate spatial IEM analysis (described above) to transform activation patterns to reconstructed spatial maps for each trial. This analysis leverages all spatial information from all voxels simultaneously to recover the stimulus representation supported by the entire neural population distributed across each retinotopic region or set of regions (Fig. 3C). To quantify stimulus representations in reconstructed spatial maps, we computed the mean map activation of pixels located within a 1.5° radius disk centered at the known position of each stimulus (matching the stimulus radius of 1.5°; see Sprague, Itthipuripat, et al., 2018; Thayer & Sprague, 2023). This provides a single value corresponding to the activation of the stimulus location for a given condition, within each retinotopic ROI. To assess the spatial selectivity of reconstructed spatial maps, we compared the mean map activation at the location of stimulus to map activation at the location opposite fixation using a 1.5° radius disk (Fig. 6 and Appendix Fig. A2).

For univariate and multivariate analyses, we computed an attention modulation index (AMI) to directly compare the impact of attending to color vs motion on stimulus representations, while accounting for the overall impact of spatial attention, similar to previous reports (McMains et al., 2007). AMI was computed as the difference in activation between the attend motion and attend color conditions, where positive values indicate a stronger response to the stimulus when attending motion (we did not normalize by the sum because, in some regions/analyses, the color- or motion-related activation was negative, making a normalized difference challenging to interpret).

In addition to the primary focused analyses on *a priori* feature-selective retinotopic regions (Figs. 4, 5), we also present all results computed for each retinotopic ROI individually (Appendix Figs. A1, A2). All figures contain data that were averaged across participants as well as individual subject means.

### Statistical analysis

Parametric statistical tests were conducted for all comparisons (repeated-measures ANOVAs and *t* tests). All statistical tests were conducted across participants after averaging all trials of each condition within individual participants. To account for possible non-normalities in our data, we generated null distributions for each test using a permutation procedure (see below) to derive p-values. Throughout the manuscript, we report statistics (F/T-scores, effect sizes) based on parametric assumptions, while p-values are derived from the shuffling procedure described above.

First, we used a one-way repeated-measures ANOVA (factor: attention condition, with 3 levels: attend fixation, attend color, attend motion) to determine whether behavioral accuracy on the fixation task depended on what was attended on each trial (Fig. 2B). We then conducted follow-up paired-samples *t* tests between all pairs of attention conditions to see which conditions differed. A paired-samples *t* test was conducted to compare attend color and attend motion RTs. The attend fixation condition was excluded for this analysis due to differences in stimulus onset (fixation target appeared briefly at least 500 ms after the dot array stimulus onset, while coherent color/motion was present throughout the dot stimulus presentation), making comparisons difficult to interpret.

When analyzing fMRI data, we focused on retinotopic regions that have previously been shown to be color- or motion-selective (Brewer et al., 2005; Huk et al., 2002; Thayer & Sprague, 2023). For univariate analyses, we computed two-way ANOVAs with attention condition (3 levels: attend fixation, color, and motion) and ROI (2 levels: color-selective hV4/VO1/VO2 and motion-selective TO1/TO2) as factors. Follow-up paired-samples *t* tests were then conducted to determine if responses differed within each ROI as a function of attention condition (Fig. 4, Appendix Fig. A1).

For multivariate neural responses, we first computed a three-way ANOVA with location (2 levels: stimulus location and opposite location), attention condition (3 levels: attend fixation, color, and motion), and ROI (2 levels: color-selective hV4/VO1/VO2 and motion-selective TO1/TO2) as factors. We then computed separate two-way ANOVAs for each relevant location (stimulus and opposite location), with attention condition (3 levels: attend fixation, color, motion) and ROI (2 levels: color-selective hV4/VO1/VO2 and motion-selective TO1/TO2) as factors. This analysis determined how task demands and feature selectivity uniquely modulated responses between regions at each location. Next, we computed the difference between stimulus and opposite locations and conducted a two-way ANOVA to determine the *spatial selectivity* of neural modulations with attention condition (3 levels: attend fixation, color, motion) and ROI (2 levels: color-selective hV4/VO1/VO2 and motion-selective TO1/TO2) as factors. To directly test whether feature-selective ROIs represent stimulus locations more strongly when participants direct attention to their preferred feature dimension, we computed a paired-samples *t* test on the AMI computed for color-selective and motion-selective ROIs (Fig. 5C, Appendix Fig. A2C). Lastly, one-sample *t* tests for univariate and multivariate AMIs were conducted for each region to determine if neural responses were feature selective (AMI different from zero). For completeness, we include data from all individual retinotopic ROIs (Appendix Fig. A5, A7) as well as the corresponding statistical analyses (Table. 1, 2). However, those results are not discussed further as there were no *a priori* hypotheses for individual regions. We note that results from individual feature-selective ROIs were weaker, but qualitatively similar to the aggregate ROI analyses.

For our shuffling procedure, we used a random number generator that was seeded with a single value for all analyses. The seed number was randomly selected using an online random number generator (https://numbergenerator.org/random-8-digit-number-generator). Within each participant, averaged data within each condition were shuffled across conditions for each participant individually, and once shuffled, the statistical test of interest was recomputed over 1000 iterations. *P* values were derived by computing the percentage of shuffled test statistics that were greater than or equal to the measured test statistic. We controlled for multiple comparisons using the false discovery rate (Benjamini & Yekutieli, 2001) across all comparisons within an analysis when necessary. Error bars are standard error, unless noted otherwise.

### Data & Code Availability

All data supporting the conclusions of this report and all associated analysis scripts are available on Open Science Framework (https://osf.io/3qrpc/?view_only=93818b413458400f9fac93dd878f09a2). To protect participant privacy, and in accordance with IRB-approved procedures, freely available data is limited to extracted timeseries for each voxel of each ROI for all scans of the study. Whole-brain ‘raw’ data will be made available from the authors upon reasonable request.

## Results

### Behavior

Participants performed a feature-selective attention task while we used fMRI to measure BOLD activation patterns across several independently defined retinotopic regions. We cued participants to covertly attend to either the color or motion of a colored-moving dot stimulus on every trial to determine which individual feature value was most prevalent. Accuracy differed across attention conditions as indicated by a permuted one-way repeated measures ANOVA, *F*(2,18) = 9.53, *p* = 0.002, 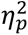 = 0.51, where accuracy was worse on attend color than attend motion trials (paired sample t-test, *t*(9) = 4.99, p < 0.001, *d_z_* = 1.58). This indicates that, despite our attempts to match difficulty, the color task was more difficult than the motion task. Notably, there was no significant difference in response times (RT; not shown) between the attend color and motion conditions (mean color and motion RT: 1.65, 1.57; SEM: 0.05, 0.06; *t*(9) = 1.42, *p* = 0.270, *d_z_* = 0.42), suggesting participants were engaged with the peripheral stimuli for a similar duration across conditions.

### Univariate neural responses

First, to determine whether the attended feature dimension impacts aggregate neural responses across independently mapped retinotopic ROIs selective for color and motion, we compared univariate neural responses averaged across all visually responsive voxels within each region, without considering their spatial selectivity with respect to the stimulus location (for individual ROI results, see Appendix Fig. A1). Hemodynamic response functions (HRFs) for each aggregate feature-selective ROI show a stimulus-related visually evoked response which differs as a function of task condition (Fig. 4A). To quantify these attentional modulations, we averaged responses 4.5-7.5 seconds after the attention cue was presented (3-6 s after stimulus onset; Fig. 4 B-C). A two-way permuted ANOVA with attention condition (attend fixation, color, and motion), and ROI (hV4/VO1/VO2 and TO1/TO2) as factors revealed a main effect of ROI (*F*(1,9) = 24.19, *p* < 0.001, 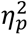 = 0.97) as well as a two-way interaction (*F*(2,18) = 54.97, *p* < 0.001, 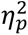 = 0.86). To determine if each region’s overall activation was modulated by attention to the peripheral stimulus as compared to attention to the fixation point, we performed follow-up permuted two-sample *t* tests within each ROI between each attend-stimulus condition and the attend fixation condition. This analysis provides insights into how directing attention to the location of the peripheral stimulus may modulate overall responses across all voxels within each region, possibly by enhancing the response to goal-relevant feature dimensions. In hV4/VO1/VO2, there was a stronger response when attending to the stimulus than when attending to fixation (fixation vs color: *t*(9) = 7.12, *p* < 0.001, *d_z_* = 2.25; fixation vs motion: *t*(9) = 4.52, *p* = 0.001, *d_z_* = 1.43), indicating that attention directed to the stimulus enhanced responses regardless of attended feature. However, TO1/TO2 showed no general modulation when attention was directed to the stimulus for either condition when compared to attending fixation (fixation vs color: *t*(9) = 2.12, *p* = 0.067, *d_z_* = 0.67; fixation vs motion: *t*(9) = 0.51, *p* = 0.598, *d_z_* = 0.16).

Next, to assess whether the specific attended feature modulates activation in each region, we computed an attention modulation index (AMI) by subtracting the neural responses between the attend motion and attend color conditions, with a positive value indicating a stronger response when attending to motion (note that this is equivalent to a paired comparison between attend-motion and attend-color conditions). Permuted one-sample *t* tests revealed that TO1/TO2 had a significant positive value (*t*(9) = 2.35, *p* = 0.024, *d_z_* = 0.74). Even though motion-selective regions showed no general modulation when attention was directed to the stimulus as compared to fixation (Fig. 4B), there was a difference in overall response as a function of attended feature dimension. In contrast, hV4/VO1/VO2 AMI based on overall activation did not significantly differ from zero (*t*(9) = 1.79, *p* = 0.061, *d_z_* = 0.56). Thus, while color regions were more active when the peripheral stimulus was relevant, their overall activation did not significantly vary based on what feature dimension was attended. Finally, a direct comparison of AMI between color- and motion-selective regions revealed a significant difference (Fig. 4C; *t*(9) = 6.07, *p* < 0.001, *d_z_* = 1.92), consistent with a region-selective modulation of overall activation levels based on attended feature.

In sum, attending to specific feature dimensions rescales the overall activation levels of feature-selective retinotopic ROIs, with regions showing greater univariate activation when their preferred feature dimension is task-relevant.

### Multivariate spatial representations

While informative, the univariate results alone cannot speak to the mechanism of attentional modulation of feature dimension maps, because univariate analyses collapsing across entire retinotopic maps are unable to discern which parts of the region—corresponding to different locations in the visual field—are modulated by task demands. Is the re-scaling observed in univariate analyses (Fig. 4) driven primarily by neural populations encoding the attended location, or instead, does this reflect a global scaling of activation across the entire ROI (Fig. 1)? To further evaluate how feature dimension maps encoded within TO1/TO2 and hV4/VO1/VO2 are modulated by task demands, we computed an inverted encoding model (IEM) for each region (Fig. 3; see Materials and Methods; Sprague et al., 2018). We used an independent “mapping” task to estimate a spatial encoding model for each voxel. The model was parameterized as a series of weights across several smooth, overlapping spatial channels (Fig. 3A). The weights from the spatial encoding model across all voxels that comprised a feature-selective ROI were then inverted which allowed us to reconstruct spatial maps based on activation profiles from the feature-selective attention task (Fig. 3B-C) averaged over 4.5-7.5 s after cue onset.

The reconstructed spatial maps (Fig. 5A) appear to have a stronger response at the aligned stimulus location than the rest of the modelled visual field (for individual ROI results, see Appendix Fig. A2). We quantified whether there was a stronger response to the stimulus by averaging map activation profiles within a 1.5° radius disk (stimulus size) at the aligned stimulus location. We could similarly assess map activation at the opposite screen location across fixation, which had no stimulus present (and was never attended; Fig. 5B). A three-way permuted ANOVA with location (stimulus and opposite), attention condition (attend fixation, color, and motion), and ROI (hV4/VO1/VO2 and TO1/TO2) revealed a three-way interaction (*F*(2,18) = 12.21, *p* < 0.001, 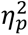 = 0.58). We conducted two-way permuted ANOVAs for each location with ROI (hV4/VO1/VO2 and TO1/TO2) and attention condition (attend fixation, color, and motion) as factors to better determine what was driving the three-way interaction. Like the univariate approach, the aim of this analysis was to determine whether motion- and color-selective regions were modulated by task demands and to determine if these regions differed in their representation depending on which feature dimension was prioritized. Responses at the stimulus location showed a main effect of attention condition (*F*(2,18) = 52, *p* < 0.001, 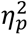 = 0.89) and ROI (*F*(1,9) = 7.31, *p* < 0.025, 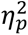 = 0.78) as well as an interaction (*F*(2,18) = 19.43, *p* < 0.001, 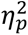 = 0.68). Follow-up permuted paired sample *t* tests indicate that color-selective regions hV4/VO1/VO2 have a stronger neural response when attending to stimulus color than when attending to fixation *t*(9) = 7.69, *p* < 0.001, *d_z_* = 2.43. There was also a stronger response when attending to stimulus motion than fixation (*t*(9) = 5.45, *p* < 0.001, *d_z_* = 1.72), consistent with spatial attention allocated to the stimulus location. Similarly, motion-selective regions TO1/TO2 responded more strongly to the stimulus when attending to either feature than when attending to fixation (fixation vs color: *t*(9) = 5.38, *p* < 0.001, *d_z_* = 1.70; fixation vs motion: *t*(9) = 5.40, *p* < 0.001, *d_z_* = 1.71), again due to spatial attention.

A two-way ANOVA at the opposite location revealed no effect of ROI (*F*(1,9) = 3.8, *p* = 0.131,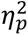 = 0.23) but did show a main effect of attention condition (*F*(2,18) = 14.56, *p* < 0.001, 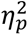 = 0.81) and an interaction between ROI and attention condition (*F*(2,18) = 3.93, *p* = 0.022, 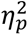 = 0.30). Follow-up permuted two-sample *t* tests reveal that there was a greater response at the opposite location in motion-selective regions when attending to fixation than either feature dimension of the stimulus (fixation vs color: *t*(9) = 4.75, *p* = 0.001, *d_z_* = 1.50; fixation vs motion: *t*(9) = 4.49, *p* = 0.001, *d_z_* = 1.42). Color-selective regions had a greater response at the opposite location when attending to fixation than attending to motion (*t*(9) = 3.19, *p* = 0.008, *d_z_* = 1.01), but there was no difference between the attend fixation and attend color conditions (*t*(9) = 2.37, *p* = 0.04, *d_z_* = 0.75, uncorrected). Thus, particularly for motion-selective regions, there are some task-dependent modulations occurring at the location opposite of the stimulus in reconstructed maps. This appears to be driven by an increased response at the opposite location when attending to fixation than when attending to either feature of the stimulus in motion regions, which is likely due to shifts of spatial attention (see Discussion).

Next, to evaluate the spatial selectivity of these modulations, we computed the difference in map activation between the stimulus and opposite locations for each attention condition (Fig. 5C). A 2-way repeated measures ANOVA with attention condition (attend fixation, color, and motion) and ROI (TO1/TO2 and hV4/VO1/VO2) as factors reveal a main effect of ROI (*F*(1,9) = 9.28, *p* = 0.018, 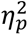 = 0.87), attention condition (*F*(2,18) = 56.92, *p* < 0.001, 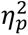 = 0.96), and an interaction between ROI and attention condition (*F*(2,18) = 12.21, *p* = 0.006, 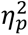 = 0.58). Follow up comparisons indicate that in color-selective regions, the stimulus location was more strongly represented when attending to a stimulus feature (fixation vs color: *t*(9) = 7.35, *p* < 0.001, *d_z_* = 2.33; fixation vs motion: *t*(9) = 7.00, *p* < 0.001, *d_z_* = 2.21). Furthermore, the stimulus location was more strongly represented when attending to color than when attending to motion (*t*(9) = 2.78, *p* = 0.019, *d_z_* = 0.88), indicating that feature-selective attention resulted in localized enhancement of the location of the stimulus representation. Motion regions also exhibit a stronger stimulus representation when attention was directed to the stimulus (attend color vs attend fixation: *t*(9) = 8.5, *p* < 0.001, *d_z_* = 2.69;attention motion vs attend fixation:*t*(9) = 9.46, *p* < 0.001, *d_z_* = 2.99). However, there was no difference between the attend color and motion conditions in the spatially selective stimulus representation (*t*(9) = 0.52, *p* = 0.610, *d_z_* = 0.16). Therefore, while TO1/TO2 is spatially selective, in the sense that spatial attention to the stimulus location locally enhances its representation, feature-selective attention results in a global shift in activation across the entire map (including the opposite location with no stimulus present). This striking and unexpected divergence between the modulatory profiles across color- and motion-selective regions suggests there may be unique spatial profiles for attentional modulation by feature relevance in color- and motion-selective regions.

Finally, we compared the AMI at the stimulus location (Fig. 5D), which indexes the difference in activation across feature-selective attention conditions, within and between color- and motion-selective ROIs. The motion-selective ROIs TO1/TO2 showed a significantly positive AMI (such that the stimulus representation was stronger during attend-motion than attend-color trials, *t*(9) = 3.34, *p* = 0.006, *d_z_* = 1.06), and the color-selective ROIs hV4/VO1/VO2 showed a significantly negative AMI (*t*(9) = 4.18, *p* < 0.001, *d_z_* = 1.32). Importantly, the AMIs derived from each set of regions significantly differed (*t*(9) = 4.5, *p* < 0.001, *d_z_* = 1.42). Altogether, this result supports a dissociable modulation of activation profiles across color- and motion-selective ROIs based on the attended feature dimension, with task-relevant feature-selective ROIs more strongly encoding stimulus locations when they are relevant for task performance.

## Discussion

The aim of the current study was to establish the spatial profile of goal-driven feature-selective attention within neural feature dimension maps (Fig. 1). We presented an identical colorful moving dot stimulus on each trial and cued participants to attend to either the color or motion of that stimulus (Fig. 2). Our design isolated the influence of observer goals because the goal-relevant feature dimension could change on each trial, but the visual input was consistent across the entire experiment. Region-level univariate methods (Fig. 4 & Appendix Fig. A1) showed unique modulations for color- and motion-selective regions: motion regions were feature-selective, but not modulated by attention directed to a peripheral stimulus as compared to fixation; color-selective regions were modulated by covert spatial attention, but this modulation was not feature-selective. However, by using an inverted encoding model (Fig. 3) to reconstruct spatial maps from both motion- and color-selective regions, we were able to show that stimulus representations in both regions were modulated by task demands and were strongest when attending to each region’s preferred feature dimension (Fig. 5 & Appendix Fig. A2). Interestingly, different spatial profiles were observed for each feature-selective region: enhancement in color regions was strongly *localized* to the stimulus relative to the opposite location, while enhancement in motion regions was not as strictly constrained to the stimulus location, consistent with some degree of global enhancement of spatial responses. Not only do these findings support the role of each region as a ‘neural feature dimension map’ (Thayer & Sprague, 2023), they extend previous work by characterizing the unique spatial profiles of attentional modulation that emerge in color- and motion-selective regions.

Previous univariate fMRI studies average across voxels with variable spatial selectivity (Kriegeskorte & Bandettini, 2007; Serences & Saproo, 2012), making it difficult to ascertain the spatial profile of attentional modulations across visual field maps encoded by feature-selective regions. However, the multivariate approach implemented here revealed that color and motion regions have unique spatial prioritization profiles. Namely, neural responses in color regions primarily differed across attention conditions at the stimulus location (Fig. 5B-D), consistent with a local enhancement of stimulus processing at task-relevant locations (Fig. 1). Motion regions also showed a stronger response at the stimulus location when motion was attended (Fig. 5B/D), along with some degree of enhancement at the opposite, task-irrelevant location (Fig. 5B/C). This is consistent with a global enhancement of motion processing across all visual field locations when motion is made task-relevant (Fig. 1), similar to previous findings for attention to a specific motion value (Saenz et al., 2002; Serences & Boynton, 2007). Characterizing these spatial profiles of prioritization due to feature-selective and spatial attention was only possible by assessing multivariate activation patterns in visual field coordinates using IEM.

Our fMRI results are consistent with participants deploying spatial attention to the stimulus location on attend-color/attend-motion trials, with a stronger observed response at the stimulus location in both regions when attending to either stimulus feature relative to fixation (Fig. 5B). If neural prioritization was solely driven by spatial attention, then multivariate responses in feature-selective regions would be equivalent between the attend color and attend motion conditions as all features at the locus of spatial attention are enhanced (Eckstein, 2011; A. Treisman, 1985; A. M. Treisman & Gelade, 1980). Rather, our AMI results indicate that activation profiles in both color- and motion-selective regions were reweighted by attention to their preferred feature dimension (Fig. 5D), suggesting that feature-selective attention acted in concert with spatial attention to sculpt activation profiles across each neural feature dimension map to support dynamic task demands (Maunsell & Treue, 2006; Scolari et al., 2014). Even though we observed clear feature preferences, it is important to highlight that these effects could have been reduced because attention can spread to irrelevant feature dimensions of an object (O’Craven et al., 1997, 1999; M. A. Schoenfeld et al., 2003, 2014), meaning that neural responses may have been enhanced even when attending to a region’s non-preferred feature dimension.

Feature-selective enhancement occurs when a feature value is relevant, resulting in an increased response from all feature-selective neural populations, regardless of spatial selectivity, and drives a spatially global enhancement that has been observed for color (McAdams & Maunsell, 2000; Saenz et al., 2002; Zhang & Luck, 2009), motion (Martínez-Trujillo & Treue, 2002; Schneider, 2011; M. Schoenfeld et al., 2007; Serences & Boynton, 2007; Treue & Trujillo, 1999), and orientation (Ester et al., 2016; McAdams & Maunsell, 2000). Interestingly, we observed more global enhancement in motion regions when attending to motion, but a local enhancement in color regions when attending to color. Notably, global prioritization through feature-selective attention is primarily observed when there are multiple stimuli at distinct locations, where one stimulus is relevant and another is irrelevant to current goals, and is strongest at the irrelevant stimulus location when the feature value matches the relevant stimulus feature (e.g., Saenz et al., 2002). If feature-selective attention modulates responses across the visual field, then all neural populations tuned to the relevant feature should have increased firing, even when there is no stimulation. For example, Serences & Boynton (2007) found evidence for successful decoding of the attended motion direction when motion was goal-relevant from voxels with no visual input.

There is evidence to suggest that color and motion have different spatial profiles when feature-selective attention is deployed. For instance, Leonard and colleagues (2015) instructed participants to identify color-defined targets in a central letter stream. When distractors matched the relevant color and were far away from the target, they weakly captured attention, but capture gradually increased as distractors were closer to the target. Similar interactions between spatial attention and feature-selective attention for color have been observed in attention and visual search tasks, where feature-specific modulations can be restricted to a single stimulus location when cued to direct spatial attention to a hemifield (Andersen et al., 2011; Berggren & Eimer, 2020; Diepen et al., 2016; Dodwell et al., 2025). This indicates that feature-selective attention for color is spatially dependent (van Es et al., 2018), diverging from the canonical characterization of spatially global prioritization of relevant features. In contrast, motion directions are almost always prioritized globally. Numerous studies have shown that when a specific motion direction is relevant, enhancement occurs throughout the visual field (Boynton et al., 2006; Mendoza et al., 2011; Treue & Trujillo, 1999), even when parametrically manipulating the distance between target and distractor motion stimuli (Liu & Mance, 2011), and when spatial attention is directed to other locations (Bosworth & Dobkins, 2002; Dobkins & Bosworth, 2001; Melcher et al., 2004; Talipski et al., 2022). Motion processing can even be improved when spatial attention is directed away from a motion stimulus (Motoyoshi et al., 2015). Furthermore, one fMRI study found increased responses in both hemifields in motion selective regions when attending to motion but only contralateral responses increases in color selective regions when attending to color (Stoppel et al., 2007). Accordingly, the evidence overall appears to point to color and motion being prioritized differently by feature- and spatial-based attention, with a greater emphasis for local enhancements of representations of color and global enhancement of representations of motion.

Our results may provide unique insight into the neural mechanisms that result in different spatial activation patterns for stimuli containing task-relevant color and motion information. The IEM approach employed here allowed for a demonstration of global modulation by feature-selective attention in motion-sensitive regions TO1/TO2, and so the absence of a similar observation for color-sensitive regions hV4/VO1/VO2 offers compelling evidence for a primarily local modulation of stimulus representations when color was task relevant. One reason why motion regions may implement global modulations while color regions locally enhance responses is that vRFs in TO1/TO2 are much larger than hV4/VO1/VO2 (Wandell & Winawer, 2015). Thus, motion regions may necessarily enhance processing across large swaths of the visual field when motion is task relevant, providing a distinct neural mechanism for the differences in spatial prioritization (McMains et al., 2007; van Es et al., 2018). This difference in encoding properties between regions may stem from the idiosyncrasies of color and motion as feature dimensions. By definition, to compute motion direction, the start and end point of an item are used to generate a direction vector (Albright & Stoner, 1995; Borst & Egelhaaf, 1989). Therefore, the global enhancement observed in the current study may be due to the specialization of TO1/TO2 in processing motion, a unique feature dimension which necessitates integration of information across distinct locations.

Notably, the current study employed different stimulus selectivity requirements than many other studies investigating attention to features. There was no requirement to prioritize a single location over others in our paradigm because only a single stimulus was ever presented. Additionally, our participants were instructed to attend to an entire feature dimension (e.g., color), rather than a specific feature value (e.g., orange). Other paradigms often include multiple stimuli and instruct participants to attend a specific feature value (Boynton et al., 2006; Mendoza et al., 2011; Treue & Trujillo, 1999; Andersen et al., 2011; Dodwell et al., 2025; van Diepen et al., 2016; Stormer & Alvarez, 2014). It is possible that the profile of attentional modulation would change across neural feature dimension maps when multiple stimuli are competing for attentional priority (Desimone & Duncan, 1995; Harrison et al., 2024; Sprague, Itthipuripat, et al., 2018), or when a specific feature value is cued as relevant.

Competing models make divergent predictions as to how task relevance of specific feature dimensions impacts activation profiles over feature dimension maps within the context of visual search. Thus, it is necessary to characterize activation profiles across the entire visual field (Fig. 3) to finely assess properties of feature dimension maps and adjudicate between these models. ‘Dimensional weighting’ models claim relevance enhances activation across entire feature dimension maps in a spatially global manner (Found & Müller, 1996; Liesefeld & Müller, 2019, 2023; Müller et al., 1995), which is consistent with our observations that attention to stimulus motion enhanced activation across the entire spatial extent of TO1/TO2. Other models claim that a spatial ‘attentional window’ restricts how much of the visual field is prioritized (Theeuwes, 2010, 2023), which is consistent with spatial activation profiles in hV4/VO1/VO2. As such, it appears aspects of both models are supported by the current findings. However, taken together, results from the current study have interesting parallels to the dimension weighting priority map model. Dimensional weighting appears to occur more weakly when color is the relevant feature dimension (Liesefeld, Liesefeld, Pollmann, et al., 2019). It has been argued that this is because color is comprised of multiple dimensions (e.g., luminance, red-green, and blue-yellow in CIE *Lab* color space). Rather than considering color to uniquely contain multiple dimensions, as motion is similarly multi-dimensional (e.g., velocity, direction, and complex motion patterns), it may be that color is unique for other reasons, including peculiarities of neural architectures supporting local processing within individual feature dimension maps. Perhaps the local modulations we observed in hV4/VO1/VO2 are the neural substrate of the finding that task-driven enhancement of color acts on a more localized portion of the visual field (Theeuwes, 2010, 2023), whereas our observation of more global enhancement in TO1/TO2 when motion is attended supports robust dimensional weighting across all spatial locations that is typically observed (Liesefeld, Liesefeld, & Müller, 2019; Liesefeld & Müller, 2019; Sauter et al., 2018).

Previous reports (McMains et al., 2007; Runeson et al., 2013) investigated how individual color- and motion-selective regions are modulated by task demands. Our findings were remarkably consistent with those found by McMains and colleagues (2007). Univariate analyses in color-selective region hV4 showed spatial selectivity, while the motion-selective region was not spatially selective. Note, that the spatial analysis conducted by McMains et al. was very coarse, as it was only able to show that the hemisphere contralateral to the stimulus had a greater response than the ipsilateral hemisphere. To assay the full spatial profile of activation relative to the stimulus location, multivariate techniques like the one implemented in the current study must be used. Importantly, our IEM approach allowed for a fully randomized stimulus position on each trial, minimizing the impact of spatial expectation effects, and eliminating the impact of any idiosyncrasies of cortical sampling of different visual field locations. Additional similarities between our results and those of McMains et al (2007) are present as there was no significant modulation based on attended feature dimension in hV4, such that there was no greater univariate response when color was goal relevant compared to motion (Appendix Fig. 1). Also converging with their results, in our study, the more anterior color-selective regions VO1 and VO2 (parallel to TEO as described in their study; Brewer et al., 2005) showed the primary evidence for color-selective attentional modulations, with statistically less-significant effects observed in hV4 (Appendix Figs. 1, 2). Based on this finding, it could be that these anterior regions are especially subject to attentional modulations. This appears to be unique to when color is goal-relevant, as hV4 qualitatively responds just as strongly, if not more strongly, to task-irrelevant but salient color stimuli than VO1/VO2 in a previous study (Thayer & Sprague, 2023). This intriguing functional distinction between extracting salient locations in posterior regions and relevant locations in anterior regions, which echoes similar sharp emergence of top-down cognitive signals in extrastriate cortex during visual working memory (Mendoza-Halliday et al., 2014), should be explored in future work spanning additional manipulations of visual stimuli and task demand conditions.

In sum, we found that feature-selective retinotopic regions in visual cortex track the location of stimuli throughout the visual field and showed a stronger response to those stimuli when each region’s preferred feature was goal-relevant. Regions had different spatial modulation profiles, with color regions locally enhancing activation at the stimulus location, and motion regions showing a global enhancement of activation. These findings emphasize the role of TO1/TO2 and hV4/VO1/VO2 as feature dimension maps and specify how they prioritize information in the visual field, which can be exploited to fully understand how we successfully navigate our visual environment.

## Acknowledgments

This work was supported by NIH R01-EY035300, an Alfred P Sloan Research Fellowship, an Nvidia Hardware Grant, and US Army Research Office Cooperative Agreement W911NF-19-2-0026 for the Institute for Collaborative Biotechnologies

## APPENDIX

**Figure A1:**
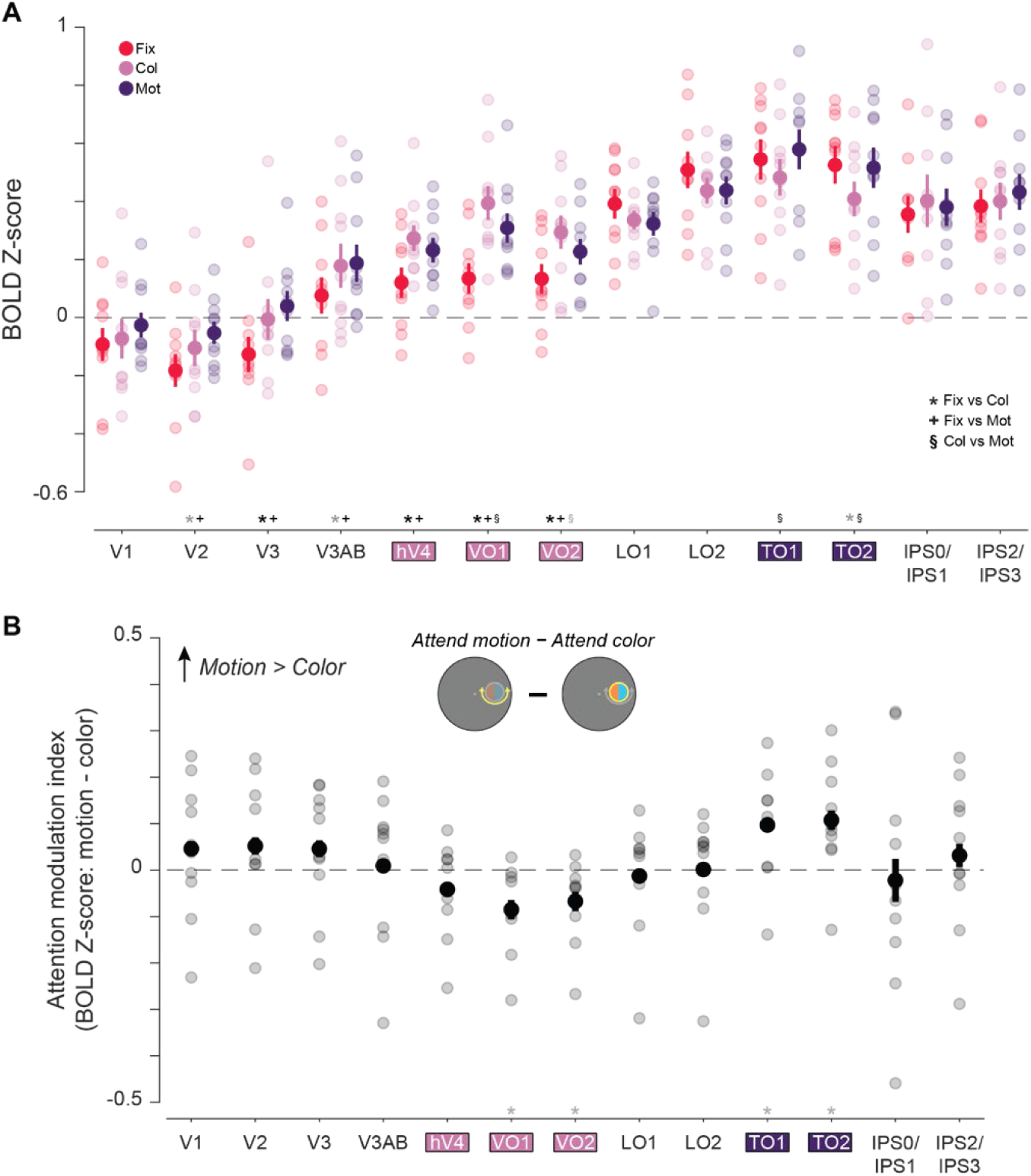
BOLD response across all voxels for individual regions of interest show selective attentional modulation based on feature preferences. **A.** Mean BOLD response for each attention condition from all individual retinotopic regions (feature-selective retinotopic regions hV4, VO1, VO2, TO1, and TO2, presented in aggregate in Fig. 4, are highlighted). Both individual motion-selective regions showed no response modulation when attending to either feature of the stimulus (color or motion) relative to when attending to fixation (all *p*’s > 0.05). Each color-selective region was modulated by task demands, with a greater region-level response when attending to the color (*p* < 0.05) or the motion (*p* < 0.05) relative to attending to fixation. **B.** Attention modulation index for each individual feature-selective region. Both motion-selective regions had a positive response that differed from zero (TO1 and TO2: *p* < 0.05, uncorrected), indicating a stronger response when attending to motion. Color-selective regions VO1 and VO2 had a negative response that differed from zero (VO1 and VO2: *p* < 0.05, uncorrected), indicating a preference for color. However, hV4 did not differ from zero (*p* > 0.05). * indicate significant differences based on permuted paired or single sample *t* tests (*p* < 0.05), corrected for multiple comparisons via FDR across 13 ROIs. Gray asterisks indicate trends, defined based on p < 0.05 without correction. Error bars indicate within-subject SEM. Faint dots indicate individual subject means. Col: color, Mot: motion, Fix: fixation.

**Figure A2:**
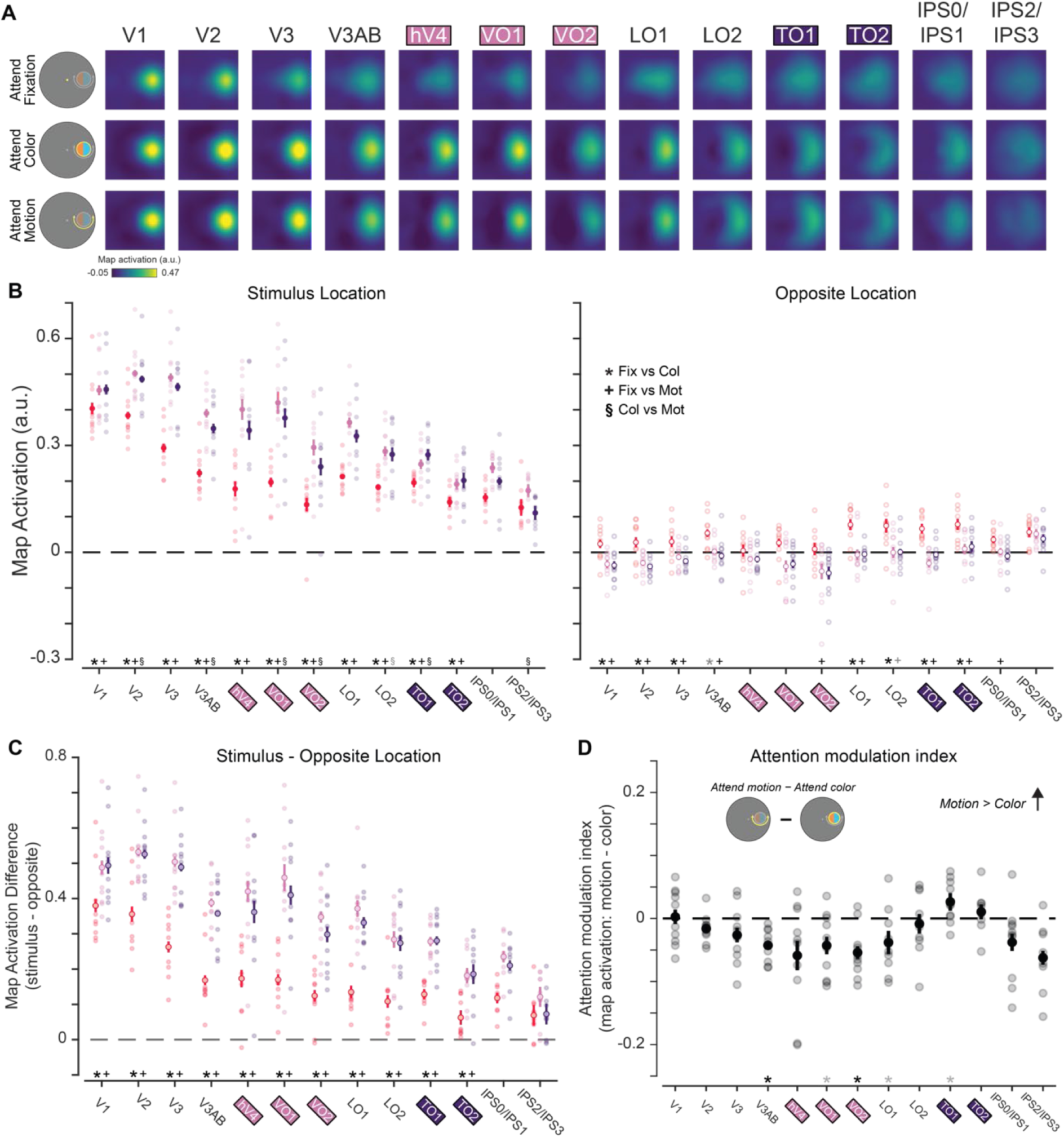
Reconstructed spatial maps for individual retinotopic ROIs. **A.** Reconstructions from all individual retinotopic ROIs, for the attend fixation, color, and motion conditions. Qualitatively, there is a strong response at the aligned location, particularly when attending to either the motion or color of the stimulus. **B.** We determined whether each region was modulated by task demands by assessing the strength of the stimulus representation for each stimulus condition as well as the opposite location. In all feature-selective regions, the neural response at the stimulus location was stronger when attending to either feature of the stimulus as compared to attend fixation trials, indicating that each region tracks task-relevance of the stimulus location. At the opposite location, responses were often strongest when attending to fixation. **C.** Difference in map activation between the stimulus and opposite locations for each stimulus condition. Responses were always greater when attending to either the color or motion of the stimulus for all regions (all *p*’s < 0.05). **D.** Difference in map activation between the attend motion and attend color conditions. Responses reliably differed from zero for TO1, VO1, and VO2, where TO1 had stronger response when attending to motion and VO1 and VO2 had stronger responses when attending to color. * Significant difference based on permuted paired-samples *t* test (*p* < 0.05), corrected for multiple comparisons via FDR. Gray asterisks indicate trends, defined as p < 0.05, uncorrected. Error bars indicate within-subject SEM. Faint dots indicate individual subject means.

**Table 1.** Statistics for univariate analyses. *P* values for univariate comparisons for individual attention conditions from zero and between specific attention conditions for each ROI. Comparisons include paired-sample *t* tests between the attend fixation, color, and motion conditions. One-sample *t* tests for individual attention conditions and AMI are also included. Even though the Color vs Motion and AMI analysis are effectively the same comparison, *p* values are slightly different due to the random shuffling procedure used to generate non-parametric distributions for statistical comparison (see statistical procedures above) and different number of comparisons when doing FDR corrections (78 for Color/Motion/Fixation comparisons; 13 for AMI). Bold denotes significance after FDR correction for all comparisons (*q* = 0.05). Significance before FDR corrections using α = 0.05 denoted with italics.

| Figures 4, 5 | Attend Fixation | Attend Color | Attend Motion | Fixation vs Color | Fixation vs Motion | Color vs Motion | AMI |
| --- | --- | --- | --- | --- | --- | --- | --- |
| V1 | 0.122 | 0.333 | 0.582 | 0.605 | 0.151 | 0.343 | 0.365 |
| V2 | **0.006** | 0.138 | 0.176 | *0.042* | **0.005** | 0.308 | 0.286 |
| V3 | 0.061 | 0.938 | 0.483 | **0.006** | **0** | 0.321 | 0.327 |
| V3ab | 0.249 | 0.041 | **0.012** | *0.032* | **0.018** | 0.848 | 0.874 |
| hV4 | 0.054 | **0** | **0** | **0.002** | **0.013** | 0.212 | 0.224 |
| VO1 | **0.023** | **0** | **0** | **0** | **0** | **0.016** | *0.013* |
| VO2 | *0.027* | **0** | **0** | **0** | **0.004** | *0.031* | *0.029* |
| LO1 | **0** | **0** | **0** | 0.234 | 0.073 | 0.781 | 0.792 |
| LO2 | **0** | **0** | **0** | 0.204 | 0.106 | 0.980 | 0.974 |
| TO1 | **0** | **0** | **0** | 0.090 | 0.234 | **0.020** | *0.024* |
| TO2 | **0** | **0** | **0** | *0.035* | 0.771 | **0.017** | *0.014* |
| TO1/TO2 | **0** | **0** | **0** | 0.067 | 0.598 | **0.014** | **0.024** |
| hV4/VO1/VO2 | **0.029** | **0** | **0** | **0** | **0** | 0.055 | 0.061 |
| IPS0/IPS1 | **0** | **0** | **0** | 0.506 | 0.617 | 0.783 | 0.788 |
| IPS2/IPS3 | **0** | **0** | **0** | 0.758 | 0.302 | 0.536 | 0.555 |

**Table 2.** Statistics for multivariate analyses. *P* values for IEM comparisons between specific attention conditions at certain locations in the visual field for each ROI. Comparisons include paired-sample *t* tests between the attend fixation, color, and motion conditions. One-sample *t* tests for AMI are also included. Even though the Color vs Motion at the stimulus location and AMI analyses are effectively the same comparison, *p* values are slightly different due to the random shuffling procedure used to generate non-parametric distributions for statistical comparison (see statistical procedures above) and different number of comparisons when doing FDR corrections (39 for Fixation/Color/Motion comparisons; 13 for AMI). Bold denotes significance after FDR correction for all comparisons (*q* = 0.05). Significance before FDR corrections using α = 0.05 denoted with italics.

| Figures 6, 7 | Stimulus Location |  |  | Opposite Location |  |  | Location Difference |  |  | AMI |
| --- | --- | --- | --- | --- | --- | --- | --- | --- | --- | --- |
|  | Fixation vs Color | Fixation vs Motion | Color vs Motion | Fixation vs Color | Fixation vs Motion | Color vs Motion | Fixation vs Color | Fixation vs Motion | Color vs Motion |  |
| V1 | **0.023** | **0.004** | 0.856 | **0** | **0** | 0.79 | **0.003** | **0** | 0.778 | 0.853 |
| V2 | **0** | **0** | **0.036** | **0.011** | **0.003** | 0.294 | **0** | **0** | 0.619 | 0.055 |
| V3 | **0** | **0** | 0.114 | **0.017** | **0** | 0.241 | **0** | **0** | 0.324 | 0.11 |
| V3ab | **0** | **0** | **0** | *0.044* | **0.006** | 0.616 | **0** | **0** | 0.185 | **0** |
| hV4 | **0** | **0** | 0.055 | 0.351 | 0.234 | 0.991 | **0** | **0** | 0.087 | 0.064 |
| VO1 | **0** | **0** | **0.023** | 0.084 | 0.057 | 0.701 | **0** | **0** | 0.099 | *0.024* |
| VO2 | **0** | **0.006** | **0.002** | 0.066 | **0.003** | 0.83 | **0** | **0.002** | 0.106 | **0** |
| LO1 | **0** | **0** | *0.048* | **0.002** | **0.003** | 0.919 | **0** | **0** | 0.157 | *0.047* |
| LO2 | **0** | **0.008** | 0.588 | **0.015** | *0.037* | 0.949 | **0** | **0.007** | 0.613 | 0.623 |
| TO1 | **0.007** | **0** | **0.042** | **0.005** | **0.002** | 0.240 | **0** | **0** | 0.912 | *0.024* |
| TO2 | **0** | **0.005** | 0.312 | **0.011** | **0.008** | 0.749 | **0** | **0** | 0.881 | 0.353 |
| TO1/TO2 | **0** | **0** | **0.011** | **0** | **0** | 0.619 | **0** | **0** | 0.610 | **0.006** |
| hV4/VO1/VO2 | **0** | **0** | **0.005** | 0.040 | **0.008** | 0.748 | **0** | **0** | **0.019** | **0** |
| IPS0/IPS1 | **0** | 0.071 | 0.080 | 0.084 | **0** | 0.555 | **0** | **0.005** | 0.280 | 0.076 |
| IPS2/IPS3 | 0.085 | 0.623 | **0.004** | 0.523 | 0.277 | 0.422 | 0.084 | 0.908 | 0.055 | **0.005** |

